# Metabolism of L-threonate, an ascorbate degradation product, requires a protein with L-threonate metabolizing domains in *Arabidopsis*

**DOI:** 10.1101/2025.05.18.654769

**Authors:** Kojiro Yamamoto, Yukino Yamashita, Tamami Hamada, Atsuko Miyagi, Hideki Murayama, Akane Hamada, Takanori Maruta

## Abstract

L-Threonate is one of the major degradation products of ascorbate in plants. While bacteria can utilize L-threonate as a sole carbon source by converting it to dihydroxyacetone phosphate, a glycolysis intermediate, through a three- or four-step metabolic pathway, the corresponding processes in plants remain uncharacterized. Remarkably, an *Arabidopsis* gene encodes a unique protein containing domains homologous to all three enzymes involved in the bacterial three-step pathway. We designated this protein as L-threonate metabolizing domains (LTD) and investigated its functional role in plant L-threonate metabolism. Despite extensive efforts, recombinant expression of LTD was unsuccessful, likely due to its large protein size. Therefore, a reverse genetic approach was employed, using *ltd* knockout *Arabidopsis* lines to explore LTD function. Under continuous dark conditions, where ascorbate degradation is facilitated, *LTD* transcription was significantly upregulated, leading to increased L-threonate dehydrogenase (LtnD) activity. Knockout lines of LTD exhibited no detectable LtnD activity under both light and dark conditions, alongside elevated levels of L-threonate compared to wild-type plants. These results indicate that LTD is essential for L-threonate metabolism in *Arabidopsis*. The *LTD* gene is highly conserved among land plants but is absent in green algae, providing a hypothesis that the rise in ascorbate concentrations during plant evolution necessitated a more active metabolism of ascorbate degradation products.

## Introduction

Ascorbate is the major water-soluble antioxidant in plants and plays a central role in maintaining intracellular redox homeostasis (Foyer and Noctor, 2005; Smirnoff, 2018). Under fluctuating environmental conditions, plants inevitably generate reactive oxygen species (ROS), which are potentially cytotoxic but also serve as crucial signaling molecules that regulate stress responses, growth, and development (Mittler et al., 2022). The antioxidant system centered around ascorbate enables plants to mitigate oxidative stress while tightly regulating ROS-mediated signaling (Maruta et al., 2016; Mittler et al., 2022). This function is particularly critical in leaves, where active photosynthesis is a primary source of ROS (Asada, 1999; Maruta, 2022). Accordingly, plants accumulate high levels of ascorbate in these tissues. To enhance ascorbate’s antioxidant capacity, plants employ ascorbate peroxidases (APX) and possess a redundant system for recycling reduced ascorbate from its oxidized forms (Maruta et al., 2024, 2016; Smirnoff, 2018). Beyond its antioxidant role, ascorbate functions as a cofactor in various metabolic processes, including phytohormone metabolism, and is involved in photosynthesis and iron acquisition (Smirnoff, 2018).

While the biosynthesis, utilization, and recycling of ascorbate have been extensively characterized (Maruta et al., 2024, 2016; Smirnoff, 2018), our understanding of its degradation remains limited, despite its importance in linking ascorbate to downstream metabolites. Ascorbate degradation is initiated by the irreversible conversion of its oxidized form, dehydroascorbate (DHA), which undergoes either oxidation or hydrolysis to produce oxalyl-L-threonate and 2,3-diketo-L-gulonate (DKG), respectively (Ford et al., 2024). These intermediates follow divergent, yet incompletely characterized, metabolic routes. In the oxalyl-L-threonate pathway, DHA (C_6_) is broken down into oxalate (C_2_) and L-threonate (C_4_) (Green and Fry, 2005). Oxalate forms calcium oxalate crystals, contributing to calcium storage and serving as a defense against herbivory (Ford et al., 2024). The DKG-dependent route appears more complex, producing a range of C_5_ sugar acids in a ROS-dependent manner. DKG oxidation may even contribute to ROS generation, potentially linking this pathway to defense responses (Kärkönen et al., 2017). In the grapevine, ascorbate is degraded into L-tartrate through a pathway involving C_4_–C_5_ bond cleavage and the intermediate L-idonate, catalyzed by L-idonate 5-dehydrogenase and 2-keto-L-gulonate reductase (DeBolt et al., 2006; Jia et al., 2019). Although L-tartrate is thought to regulate fruit cell pH, its physiological role remains poorly understood (Burbidge et al., 2021).

Despite being a product of the oxalyl-L-threonate pathway, the metabolism and function of L-threonate in plants remain largely unexplored. Nevertheless, L-threonate accumulation is observed under stress conditions. For instance, *Arabidopsis thaliana* catalase-deficient mutants (*cat2*) exhibit elevated L-threonate levels (Noctor et al., 2015), likely due to enhanced ascorbate oxidation and degradation. Similarly, high-light exposure increases L-threonate levels in *Arabidopsis* wild-type plants but not in the ascorbate-deficient mutant (*vitamin c defective 2, vtc2*) (Terai et al., 2020). In multiple mutants defective in ascorbate recycling, inhibition of ascorbate accumulation after high light is accompanied by a pronounced increase in L-threonate levels (Terai et al., 2020). These findings suggest that L-threonate accumulation results from accelerated ascorbate degradation when recycling is impaired. However, whether L-threonate accumulation has physiological significance or merely reflects stress-induced catabolism remains unclear.

In bacteria, L-threonate can serve as a sole carbon source by being converted into dihydroxyacetone phosphate (DHAP), an intermediate in glycolysis. The initial four-step bacterial pathway involves L-threonate dehydrogenase (LtnD), 2-oxo-tetronate isomerase (OtnI), 3-oxo-tetronate kinase (3-OtnK), and 3-oxo-tetronate 4-phosphate decarboxylase (3-OtnC) (Zhang et al., 2016) (**Fig. 1**). In this pathway, L-threonate is first oxidized by LtnD to produce 2-oxo-L-threonate. This intermediate then undergoes an isomerization reaction catalyzed by OtnI to yield 3-oxo-L-threonate. Subsequently, 3-oxo-L-threonate is phosphorylated by 3-OtnK to form 3-oxo-4-phospho-L-threonate, which is finally converted into DHAP through a decarboxylation reaction catalyzed by 3-OtnC. Deletion of these genes abolishes the ability to utilize L-threonate as a carbon source (Zhang et al., 2016). A three-step bypass pathway, which omits OtnI, was recently identified, in which 2-oxo-L-threonate generated by LtnD is directly converted to DHAP via 2-OtnK and 2-OtnC (Guo et al., 2024) (**Fig. 1**).

**Figure 1.**
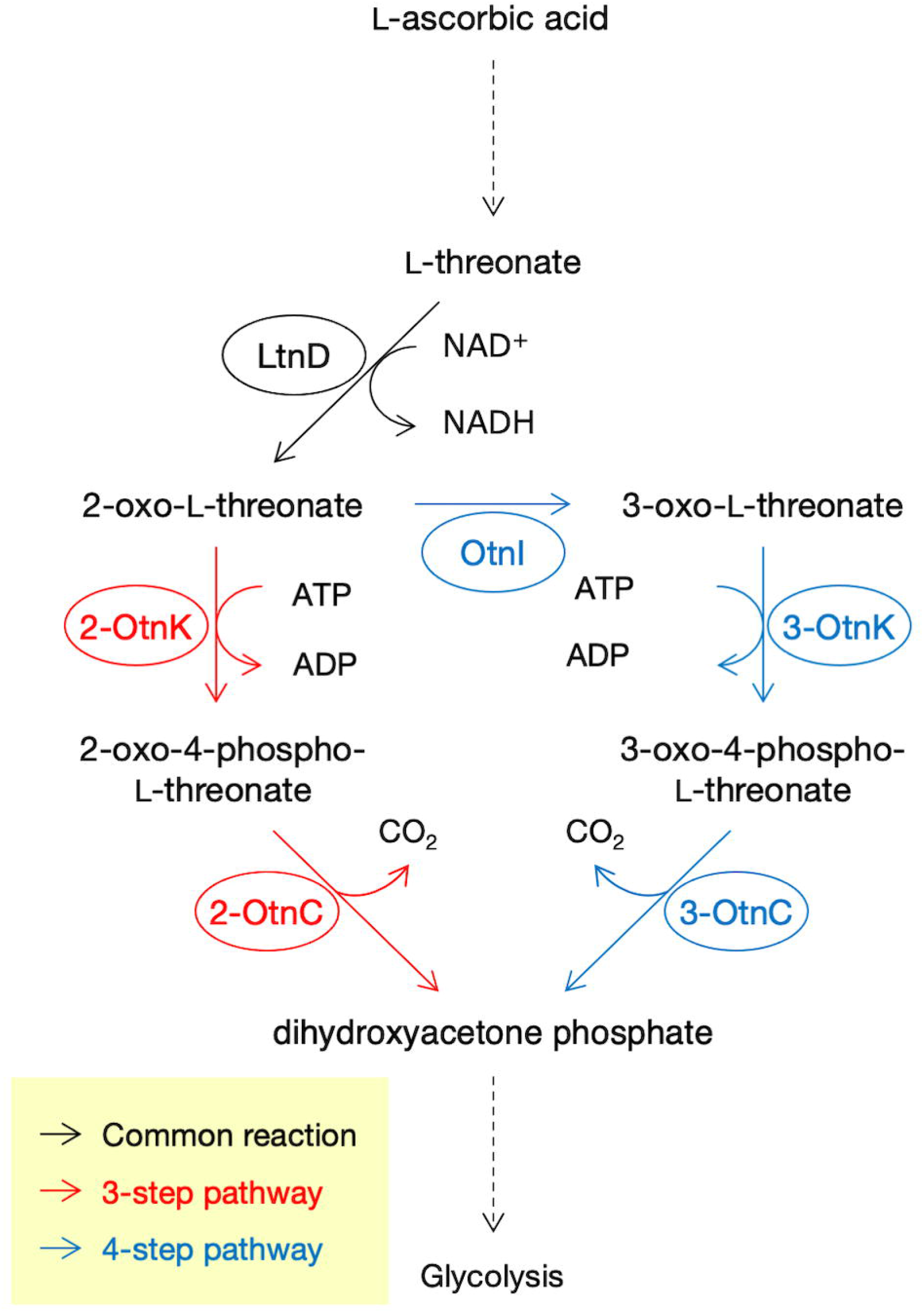
Bacterial pathways for L-threonate catabolism Schematic representation of the bacterial L-threonate catabolic pathways. In the four-step pathway, L-threonate is oxidized by L-threonate dehydrogenase (LtnD) to form 2-oxo-L-threonate, which is then isomerized by 2-oxo-tetronate isomerase (OtnI) to generate 3-oxo-L-threonate. This compound is phosphorylated by 3-oxo-tetronate kinase (3-OtnK) to produce 3-oxo-4-phospho-L-threonate, which is subsequently converted into dihydroxyacetone phosphate (DHAP) by 3-oxo-tetronate 4-phosphate decarboxylase (3-OtnC). In the three-step bypass pathway, 2-oxo-L-threonate is directly converted to DHAP via 2-OtnK and 2-OtnC, bypassing the OtnI-mediated isomerization step.

Of particular interest, an uncharacterized *Arabidopsis* gene, *AT1G18270*, encodes a single polypeptide containing domains homologous to bacterial LtnD, OtnK, and OtnC. This suggests that *AT1G18270* may be a key, yet previously uncharacterized, component of L-threonate metabolism in plants. In this study, we refer to this gene as L-threonate metabolizing domains (*LTD*) and investigate its function. Our findings reveal that LTD is indispensable for L-threonate metabolism in *Arabidopsis*.

## Results

### Domain structure of LTD and its distribution in the plant kingdom

The *Arabidopsis LTD* gene encodes a unique protein containing domains homologous to three bacterial enzymes involved in the three-step L-threonate catabolic pathway. The domain architecture of *Arabidopsis* LTD is shown in **Fig. 2**. From the N-terminus, two consecutive NAD-binding domains, NAD_binding_2 and NAD_binding_11, are present, a feature conserved in LtnD from *Escherichia coli* (four-step pathway) and *Arthrobacter* sp. ZBG10 (three-step pathway) (Guo et al., 2024; Zhang et al., 2016). This tandem arrangement of NAD-binding domains appears twice in succession. For convenience, we refer to the first as LtnD1 and the second as LtnD2. *Arabidopsis* LtnD1 and LtnD2 share 26.2–38.7% identity and 44.6– 60.7% similarity with the corresponding enzymes from *E. coli* and *Arthrobacter* (**Supplementary Fig. S1**).

**Figure 2.**
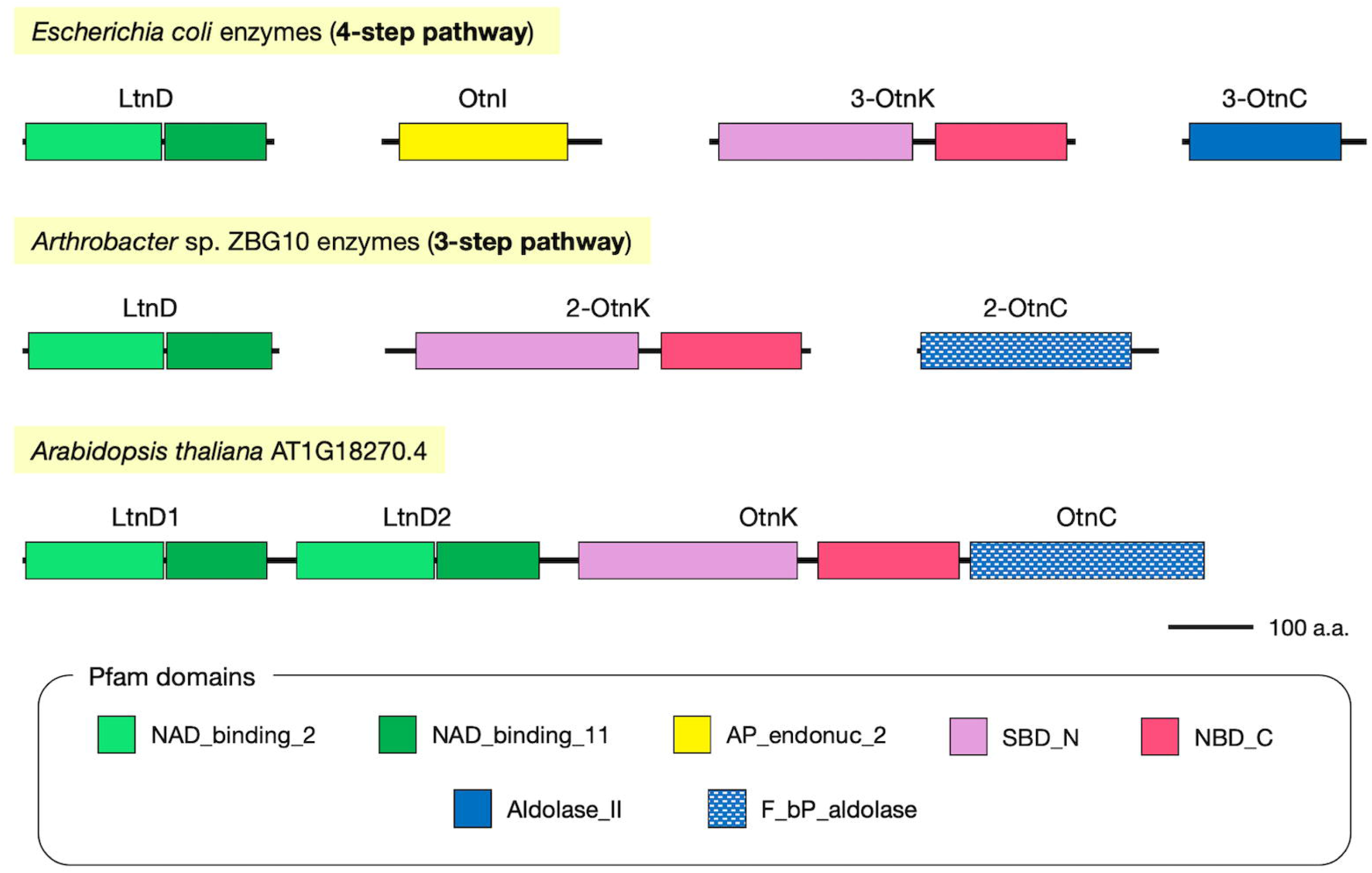
Domain architectures of bacterial and Arabidopsis enzymes involved in L-threonate catabolism Schematic comparison of the domain structures of enzymes involved in L-threonate catabolism in *Escherichia coli* (four-step pathway), *Arthrobacter* sp. ZBG10 (three-step pathway), and *Arabidopsis thaliana* (AT1G18270.4, LTD). Bacterial enzymes are shown individually, while Arabidopsis LTD integrates all three functional modules (LtnD, OtnK, and OtnC) into a single polypeptide. LTD contains two tandem NAD-binding regions (LtnD1 and LtnD2), followed by SBD_N and NBD_C domains (OtnK-like), and a C-terminal F_bP_aldolase domain (OtnC-like). Domain annotations are based on Pfam predictions.

Following the LtnD-like domains, the protein contains two additional domains: SBD_N and NBD_C, arranged sequentially (**Fig. 2**). These are homologous to 3-OtnK from *E. coli* and 2-OtnK from *Arthrobacter* sp. ZBG10. The identity and similarity of the *Arabidopsis* OtnK domain to *E. coli* 3-OtnK are 18.5% and 32.7%, respectively, while the values are higher when compared with *Arthrobacter* 2-OtnK (32.0% and 46.7%, respectively) (**Supplementary Fig. S1**).

Finally, the C-terminal region of *Arabidopsis* LTD contains an F_bP_aldolase domain, which is similar to the 2-OtnC domain of *Arthrobacter*, but distinct from the Aldolase_II domain of *E. coli* 3-OtnC (**Fig. 2**). Indeed, the OtnC domain of *Arabidopsis* LTD shows negligible identity and similarity to *E. coli* 3-OtnC but shares 33.2% identity and 47.7% similarity with *Arthrobacter* 2-OtnC (**Supplementary Fig. S1**). These findings suggest that *Arabidopsis* LTD may functionally encode the entire three-step L-threonate metabolic pathway found in bacteria, such as *Arthrobacter*. This protein lacks organellar targeting signals and transmembrane domains, suggesting that it localizes to the cytosol.

To investigate the evolutionary conservation of *LTD* in plants, we searched for *LTD* homologs in green algae, bryophytes, liverworts, ferns, and angiosperms. The *LTD* gene was found in all examined land plant genomes (such as *Marchantia polymorpha, Physcomitrium patens, Selaginella moellendorffii*, and *Oryza sativa*), but not in green algae (*Chlamydomonas reinhardtii, Dunaliella salina, Micromonas* sp RCC299, *Ostreococcus lucimarinus*, and *Volvox carteri*). Cyanobacteria are known to possess the three-step L-threonate catabolic pathway (Guo et al., 2024), making the absence of LTD or L-threonate metabolizing enzymes in green algae intriguing. However, the evolutionary significance of this loss remains unclear. The domain architectures of LTD homologs from various land plants are shown in **Supplementary Fig. S2**. Like *Arabidopsis*, LTD homologs in bryophytes, ferns, and rice possess two LtnD-like domains, one 2-OtnK-like domain, and one 2-OtnC-like domain. These results indicate that LTD is a highly conserved and previously uncharacterized gene family in land plants.

### Recombinant expression of LTD and attempts to detect LtnD activity

To investigate whether *Arabidopsis LTD* encodes the three-step pathway found in bacteria, we attempted to produce recombinant LTD protein. First, we cloned the full-length LTD coding sequence and attempted to express it in *E. coli*. Despite testing multiple expression vectors, host strains, and cultivation/induction conditions, we were unable to achieve high-level expression of full-length LTD in *E. coli*. This difficulty was likely due to the large size of the LTD protein. We next attempted to express the region encoding both LtnD1 and LtnD2, but this also failed. Finally, we expressed LtnD1 and LtnD2 individually, and only LtnD1 was successfully represented in the soluble fraction of *E. coli* only when fused to a trigger factor (TF) tag, which has chaperone activity (**Supplementary Fig. S3**). After affinity purification using an N-terminal His-tag, we attempted to remove the TF tag using protease digestion, but the tag could not be separated from LtnD1. Therefore, we used the purified LtnD1-TF fusion protein for activity assays (**Supplementary Fig. S3**). We optimized the assay conditions based on previous studies, but unfortunately, we were unable to detect LtnD activity from LtnD1-TF. Possible reasons for this include improper folding of LtnD1 when expressed alone, inactivation during expression or purification, or interference by the TF tag.

### LtnD activity in Arabidopsis is LTD-dependent

Since the use of recombinant enzyme was technically challenging at this stage, we next attempted to detect LtnD activity using *Arabidopsis* shoots. We focused on LtnD because the substrates for 2-OtnK and 2-OtnC are not commercially available. If the detected LtnD activity proved to be LTD-dependent, it would provide evidence supporting the involvement of LTD in L-threonate metabolism. To this end, we obtained two independent T-DNA insertion lines for LTD and isolated homozygous mutants. Under standard *Arabidopsis* growth conditions, the *ltd* mutants displayed no visible differences from the wild type (**Supplementary Fig. S4**).

Bacterial LtnD uses Mg^2+^ as a cofactor (Guo et al., 2024; Zhang et al., 2016). We first homogenized *Arabidopsis* wild-type shoots, prepared soluble protein extracts, and measured Mg^2+^-dependent L-threonate dehydrogenase activity by monitoring NADH production spectrophotometrically. However, no activity was detected. We also tested Mn^2+^ and Zn^2+^ as alternative cofactors, as well as reactions without added cofactors, but again, no activity was observed. Further optimization of buffer composition, pH, and temperature also failed to yield detectable activity.

The difficulty in detecting LtnD activity was likely due to low intrinsic enzyme activity and/or low LTD expression under standard growth conditions. As L-threonate metabolism is presumably more important under conditions in which ascorbate degradation is enhanced, we examined *LTD* transcript levels under prolonged darkness, a condition known to promote ascorbate breakdown (Truffault et al., 2017; Yabuta et al., 2007). Two-week-old wild-type *Arabidopsis* plants were placed in the dark for 72 hours. Ascorbate levels in shoots decreased markedly over time (**Fig. 3a**), and *LTD* transcript levels increased six-fold after 24 hours of dark treatment and remained high thereafter (**Fig. 3b**). We then attempted to detect LtnD activity using samples prepared under these conditions. Soluble extracts from shoots before and after dark treatment were concentrated using ultrafiltration columns, and similar enzyme assays were performed. However, even in concentrated samples, no LtnD activity was detected. Moreover, the addition of cofactors to the reaction mixture with concentrated samples caused protein aggregation, particularly in the presence of Zn^2+^. Therefore, we abandoned the standard spectrophotometric assay based on NADH detection.

**Figure 3.**
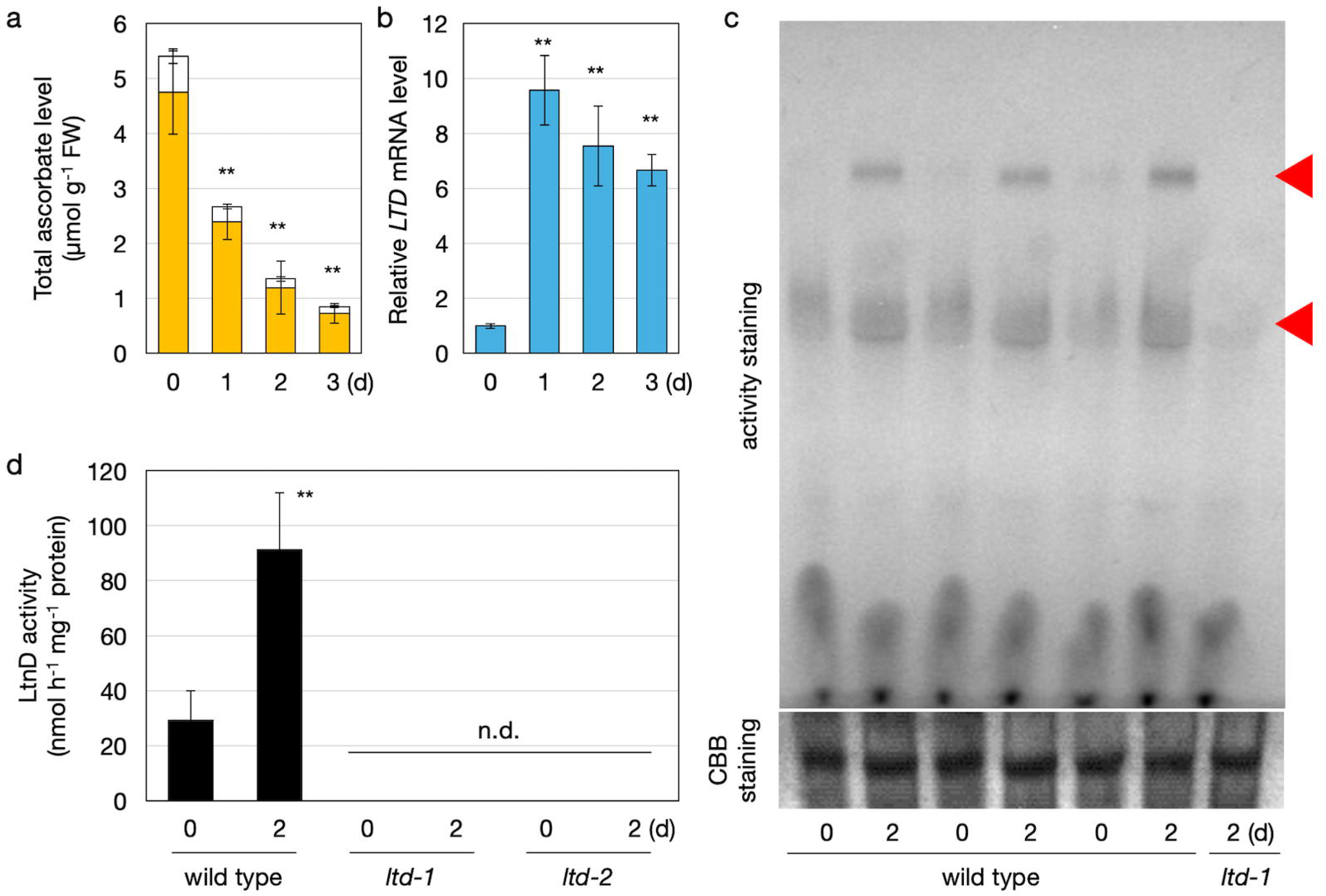
*LTD* expression and LtnD activity in Arabidopsis wild type and *ltd* mutants *Arabidopsis thaliana* wild-type (Col-0), *ltd-1*, and *ltd-2* plants were grown on half-strength MS medium without sucrose for two weeks and then transferred to dark conditions. Shoots were harvested and used for analysis. (a) Total ascorbate content (sum of ASC and DHA). (b) Transcriptional levels of LTD. (a, b) Data are presented as mean ± SD of three or four biological replicates. Statistical differences between samples before and after dark treatment were determined by one-way ANOVA followed by Dunnett’s post hoc test. ^**^*P* < 0.01, compared to values at 0 d. (c) Concentrated protein extracts were separated by 3–14% gradient native PAGE and subjected to activity staining for LtnD. The two red triangles in the upper panel indicate activity signals. Wild-type samples before and after 2 days of dark treatment (three biological replicates) were run on the same gel. The *ltd-1* sample after 2 days of dark treatment was used as a negative control. The lower panel shows CBB staining as a loading control. (d) LtnD activity. No activity was detected in *ltd-1* or *ltd-2*. Data are presented as mean ± SD of four biological replicates. Statistical differences between samples before and after dark treatment in the wild type were determined by Student’s *t*-test. ^**^*P* < 0.01. n.d., not detected.

We next tested an activity staining method. Concentrated crude extracts were subjected to native PAGE and incubated with a reaction mixture containing L-threonate, NAD^+^, phenazine methosulfate (PMS), and nitroblue tetrazolium (NBT). In this system, NADH produced by LtnD activity reduces PMS, which in turn reduces NBT to form a blue-purple formazan precipitate. Using this method, we successfully detected LtnD activity from *Arabidopsis* shoots. Two distinct activity bands were observed, both of which were enhanced following dark treatment (**Fig. 3c**), which reflected the upregulation of *LTD* transcription (**Fig. 3b**). Although further analysis is needed, the higher-molecular-weight band might be derived from a dimeric form of LTD. Importantly, neither of these signals was detected in the dark-treated *ltd* mutants (**Fig. 3c**), indicating that both bands were LTD-dependent.

Furthermore, by quantifying the formazan production spectrophotometrically, we were able to determine the LtnD activity level. Cofactors were not added in this assay (as well as in the activity staining method), as higher activity was detected in the absence of added cofactors than when Mg^2+^ was supplied. This suggests that *Arabidopsis* LtnD may bind tightly to its cofactors, making external addition unnecessary for activity detection. The LtnD activity in two-week-old shoots (4 hours after light exposure) was measured at 29.1 nmol h^-1^ mg^-1^ protein, and this value increased approximately 3.1-fold after two days in the dark (**Fig. 3d**). No LtnD activity was detected in either *ltd* mutant line by any of the methods used, confirming that detectable LtnD activity in *Arabidopsis* is LTD-dependent.

We then used this established method to test dehydrogenase activity toward other C_4_ sugar acids, including D-threonate, L-erythronate, D-erythronate, and L-tartrate, all of which are known or potential substrates of bacterial LtnDs. Among these, only D-erythronate dehydrogenase activity was detected in *Arabidopsis* shoot extracts, but this activity was not dependent on LTD (**Supplementary Fig. S5**). No or negligible dehydrogenase activity was detected toward the other tested compounds (**Supplementary Fig. S5**).

### LTD is essential for L-threonate metabolism

To determine whether LTD is required for L-threonate metabolism, we quantified L-threonate levels in shoots of wild-type and *ltd* lines before and after dark treatment. Before dark treatment, L-threonate levels tended to be higher in *ltd* mutants than in the wild type (**Fig. 4**), with a statistically significant difference between wild type and *ltd-1*. Interestingly, while L-threonate levels did not increase after dark treatment in wild-type plants, both *ltd* lines exhibited marked L-threonate accumulation (**Fig. 4**). These results demonstrate that LTD is essential for L-threonate metabolism. In wild-type plants, the dark-induced increase in LtnD activity likely suppressed L-threonate accumulation completely.

**Figure 4.**
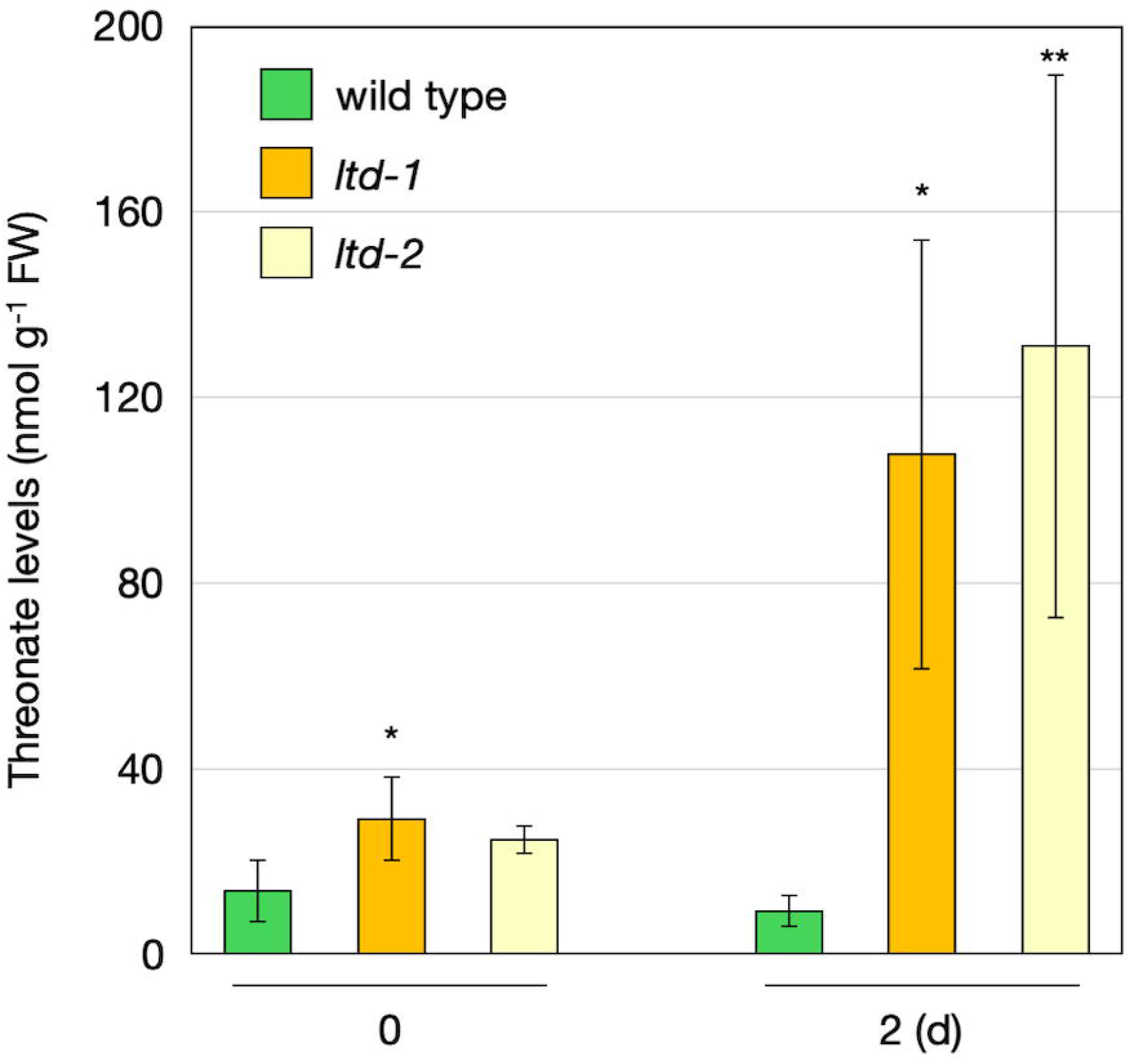
L-threonate levels in Arabidopsis wild type and *ltd* mutants in the dark *Arabidopsis thaliana* wild-type (Col-0), *ltd-1*, and *ltd-2* plants were grown on half-strength MS medium without sucrose for two weeks and then transferred to dark conditions. Shoots were harvested and used for L-threonate quantification. Data are presented as mean ± SD of four biological replicates. Statistical differences among genotypes were determined by one-way ANOVA followed by Dunnett’s post hoc test: ^*^*P* < 0.05, ^**^*P* < 0.01, compared to the wild type.

### Stress responses of LTD transcription and LtnD activity

As ascorbate degradation is enhanced under oxidative stress and high-light conditions (Noctor et al., 2015; Terai et al., 2020), we next examined the transcriptional response of *LTD* to oxidative stress. Two-week-old wild-type plants were treated for 6 hours with either high light (1,500 µmol photons m^-2^ s^-1^) or 25 µM paraquat. *LTD* transcript levels increased approximately two-fold in response to paraquat, but did not change under high light (**Fig. 5a**). The lack of response to high light may have been due to the relatively weaker oxidative stress compared to paraquat treatment. However, LtnD activity did not increase in response to paraquat, but rather decreased (**Fig. 5b**).

**Figure 5.**
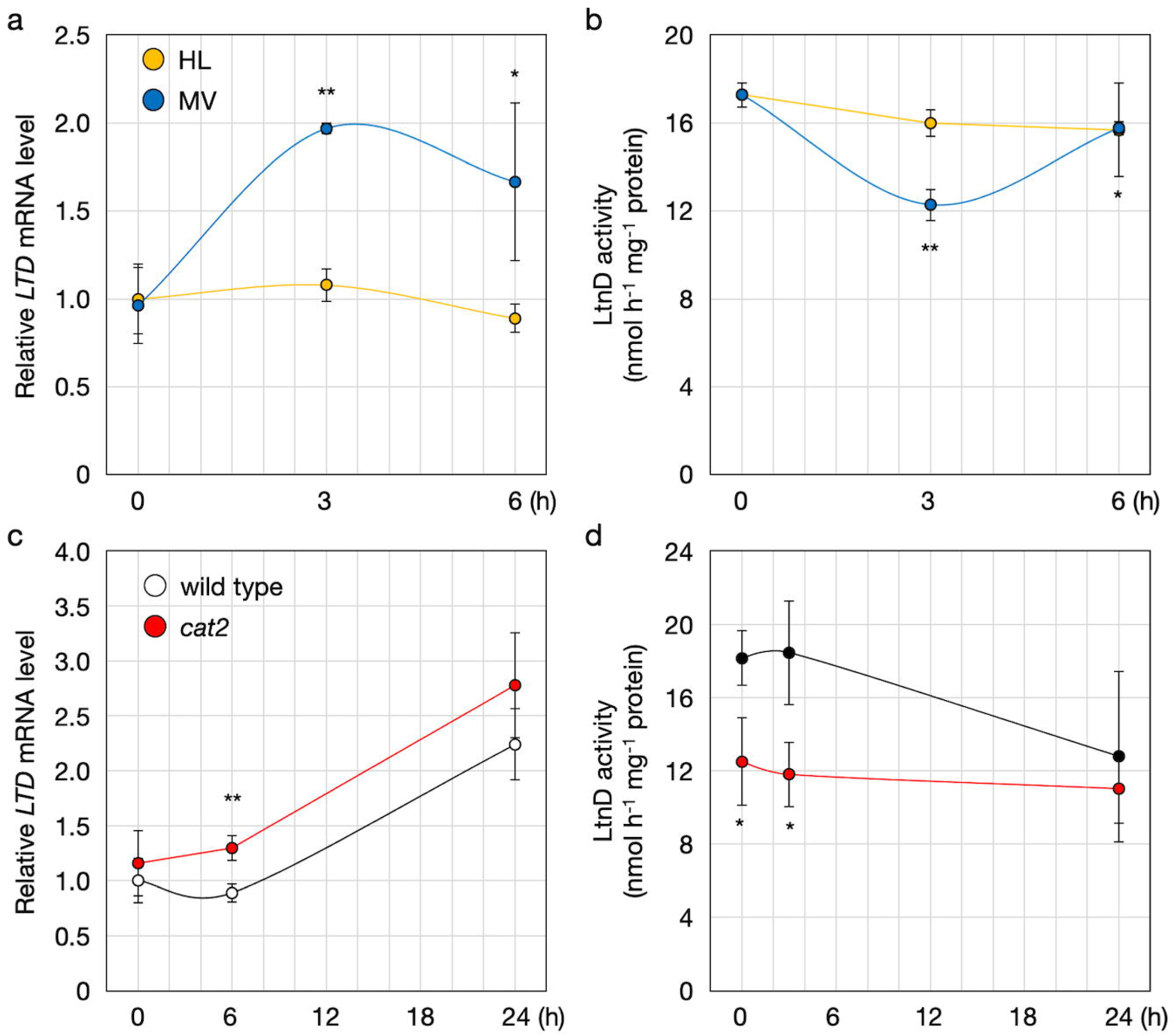
*LTD* expression and LtnD activity in Arabidopsis under photooxidative stress (a, b) *Arabidopsis thaliana* wild-type (Col-0) plants were grown on half-strength MS medium without sucrose for two weeks and then subjected to high-light stress (HL, 1,500 µmol photons m^-2^ s^-1^) or sprayed with 25 µM methyl viologen (MV) for 6 hours. Shoots were harvested for analysis of *LTD* transcript levels (a) and LtnD activity (b). Data are presented as mean ± SD of three or four biological replicates. Statistical differences between time points were determined by one-way ANOVA followed by Dunnett’s post hoc test. ^*^*P* < 0.05, ^**^*P* < 0.01, compared to 0 h. (c, d) Wild-type and *cat2* plants were grown under the same conditions and subjected to HL stress for 24 hours. Shoots were harvested for analysis of *LTD* transcript levels (c) and LtnD activity (d). Statistical differences between genotypes were determined by Student’s *t*-test. ^*^*P* < 0.05, ^**^*P* < 0.01.

For a more detailed analysis, we used the oxidative stress model mutant *cat2*. Wild-type and *cat2* plants were exposed to high light, and *LTD* transcript levels and LtnD activity were examined after short (3 h) and long (24 h) exposure. As observed previously, *LTD* transcription was unresponsive to short-term high light but was induced after 24 hours. Interestingly, this induction was slightly greater in *cat2* than in the wild type (**Fig. 5c**). However, LtnD activity did not increase in either genotype after light exposure and was even lower in *cat2* than in the wild type (**Fig. 5d**). Thus, although *LTD* transcription responded to strong or prolonged oxidative stress, LtnD enzymatic activity tended to decrease under these conditions.

## Discussion

L-threonate is one of the degradation products of ascorbate. The ascorbate degradation pathway in plants is highly complex, and not all degraded ascorbate is converted into L-threonate (Ford et al., 2024; Smirnoff, 2018). Nevertheless, considering the exceptionally high concentrations of ascorbate in plant tissues (Gest et al., 2013), it is physiologically plausible that large amounts of L-threonate are generated under conditions where ascorbate degradation is enhanced, such as prolonged darkness or severe oxidative stress (Noctor et al., 2015; Terai et al., 2020; Truffault et al., 2017). To understand the physiological relevance and consequences of L-threonate generation, it is important to elucidate its metabolism in plants. In this study, we focused on *Arabidopsis* LTD, which contains domains homologous to bacterial L-threonate metabolic enzymes, and investigated its function.

*Arabidopsis* LTD is a unique protein containing domains homologous to LtnD, OtnK, and OtnC, all of which are involved in the bacterial L-threonate catabolic pathway (Guo et al., 2024; Zhang et al., 2016). The main distinction between the three-step and four-step bacterial pathways lies in the presence of OtnI, which is required for the isomerization of 2-oxo-L-threonate only in the four-step pathway (**Fig. 1**). Additionally, the domain structure of OtnC differs between the two pathways: 2-OtnC in the three-step pathway contains an F_bP_aldolase domain, whereas 3-OtnC in the four-step pathway contains an Aldolase_II domain (Guo et al., 2024) (**Fig. 2**). *Arabidopsis* LTD lacks an OtnI-like domain and contains an F_bP_aldolase domain (**Fig. 2**). We performed a BLASTP search using the amino acid sequence of *E. coli* OtnI as a query, but no sequences homologous to OtnI were found in *Arabidopsis*. These findings suggest that LTD includes all the domains homologous to the bacterial three-step pathway. LTD homologs were found in all examined land plants, including bryophytes, ferns, and flowering plants (**Supplementary Fig. S2**). Their domain structures were identical to that of *Arabidopsis* LTD, suggesting that the three-step pathway is highly conserved in land plants.

Despite considerable effort, we were unable to produce full-length recombinant LTD protein. LtnD1 was the only domain that could be successfully expressed as a soluble protein, and only when fused to a chaperone tag (**Supplementary Fig. S3**). However, this fusion protein exhibited no detectable LtnD activity, possibly due to misfolding, inactivation during purification, or interference from the tag. Although further biochemical studies are required, our reverse genetic approach provided clear evidence for the essential role of LTD in L-threonate metabolism. In wild-type plants, dark-induced ascorbate degradation was accompanied by increased *LTD* transcription and LtnD activity (**Fig. 3**). In contrast, LtnD activity was undetectable in two independent *ltd* lines. Notably, L-threonate levels were significantly higher in *ltd* mutants than in the wild type, particularly under dark conditions (**Fig. 4**). These findings strongly support the requirement of LTD for L-threonate metabolism. The lack of L-threonate accumulation in wild-type plants under dark conditions suggests that the increase in LtnD activity enabled its rapid degradation (**Fig. 4**). If LTD indeed performs the full three-step bacterial pathway as a single protein, then L-threonate generated from ascorbate under dark conditions could be converted to DHAP and enter glycolysis. Although this hypothesis requires further enzymatic validation, it raises the possibility that LTD is involved in recycling carbon from ascorbate.

L-threonate accumulation occurs under oxidative stress (Noctor et al., 2015; Terai et al., 2020). Since L-threonate accumulation under high light is suppressed in ascorbate-deficient *vtc2* mutants (Terai et al., 2020), the accumulated L-threonate likely originates from ascorbate. Although *LTD* transcription was induced under oxidative stress, including paraquat treatment, prolonged high light, and in *cat2*, LtnD activity did not increase and instead tended to decrease (**Fig. 5**). The mismatch between transcriptional induction and enzyme activity suggests the existence of post-transcriptional or post-translational regulation of LTD. Suppression of LtnD activity under oxidative stress may permit L-threonate accumulation in such conditions as previously observed (Noctor et al., 2015; Terai et al., 2020). While *ltd* mutants showed no apparent phenotype under standard growth conditions (**Supplementary Fig. S4**), a comparison of stress sensitivities may help uncover the physiological functions or effects of L-threonate accumulation under stress conditions.

The enzymes constituting the bacterial three-step pathway are also present in cyanobacteria (Guo et al., 2024). Although LTD is highly conserved in land plants, it is absent from green algae, possibly due to secondary loss (**Supplementary Fig. S2**). The reason for this is unclear, but green algae contain much lower ascorbate levels than land plants (Gest et al., 2013; Maruta et al., 2024). Thus, the small amounts of L-threonate produced in green algae may be degraded by alternative mechanisms or excreted. In contrast, land plants may generate substantial levels of L-threonate and, being unable to excrete it, may have needed to reacquire a metabolic pathway for its degradation.

In summary, our study identifies LTD as a multi-domain protein structurally and functionally related to the bacterial three-step L-threonate degradation pathway. We show that LTD is indispensable for LtnD activity and L-threonate catabolism in planta. These findings provide a foundation for future biochemical, physiological, and genetic studies to elucidate the full picture of ascorbate degradation and uncover the physiological significance of L-threonate metabolism in plants.

## Materials and Methods

### Plant materials and growth conditions

*Arabidopsis thaliana* ecotype Col-0 was used as the wild-type plant. The T-DNA insertion mutants *ltd-1* (SALK_110477C), *ltd-2* (SALK_044797C), and *cat2* (SALK_057998) were obtained from the Arabidopsis Biological Resource Center (ABRC). Wild-type and mutant seeds were sown on half-strength Murashige and Skoog (MS) medium without sucrose and stratified in the dark at 4°C for 2–3 days. Plants were grown under a 16-hour light/8-hour dark photoperiod at 22°C/20°C, respectively, with a light intensity of 100 µmol photons m^-2^ s^-1^ (LH-240S, NK System) for two weeks. For dark treatment, plates were wrapped with two layers of aluminum foil and incubated under the same growth conditions. For photooxidative stress experiments, 2-week-old plants were either exposed to high light (1,500 µmol photons m^-2^ s^-1^) or sprayed with 25 µM methyl viologen (MV) solution containing 0.1% Tween 20 (v/v), followed by incubation under standard light conditions. For growth comparison (Supplemental Figure S4), seeds were sown in Jiffy-7 peat pellets, stratified at 4°C in the dark for 2–3 days, and grown under the same conditions for three weeks.

### Cloning and expression of LTD in *Escherichia coli*

Total RNA was extracted from wild-type shoots using RNAiso Plus (Takara) and treated with DNase to eliminate genomic DNA contamination. First-strand cDNA was synthesized using ReverTra Ace reverse transcriptase (Toyobo) with an oligo(dT) primer. Full-length *LTD* cDNA, as well as domain-specific cDNAs encoding LtnD1 and/or LtnD2, were amplified from the first-strand cDNA using primers listed in **Supplemental Table S1**. The resulting PCR products were cloned into the *E. coli* expression vector pCold-II or pCold TF (which encodes a trigger factor [TF] tag) using In-Fusion cloning. The *E. coli* strain BL21 (DE3) pLysS was used as the host for recombinant protein expression. Transformed cells were cultured at 37°C in 400 mL Luria-Bertani (LB) medium supplemented with 50 µg mL^-1^ ampicillin and 34 µg mL^-1^ chloramphenicol. When the culture reached an optical density of 0.4–0.5 at 600 nm, protein expression was induced by adding 0.5 mM isopropyl β-D-1-thiogalactopyranoside (IPTG), followed by incubation at 15°C overnight. Cells were collected by centrifugation and stored at −20°C until use.

For protein extraction, frozen cells were resuspended in 20 mM sodium phosphate buffer (pH 7.0) containing 300 mM NaCl and lysed by sonication. Recombinant proteins were purified using a TALON Metal Affinity Resin column (Clontech) and concentrated using a 30-kDa cutoff ultrafiltration device (Amicon® Ultra-4, Millipore). We attempted to cleave the TF tag from the recombinant protein using HRV 3C Protease (Takara); however, the tag could not be separated from the target protein, likely due to incomplete cleavage or tight association with the protein.

### Semi-quantitative and quantitative reverse transcription-polymerase chain reaction

Semi-quantitative reverse transcription-polymerase chain reaction (RT-PCR) was performed according to Terai et al. (2020). Quantitative RT-PCR (qPCR) was performed as described by Kameoka et al. (2021). All the primers used are listed in **Supplementary Table S1**.

### Measurements of ascorbate and threonate

Ascorbate was quantified using an ultra-fast liquid chromatography (UFLC) system (Shimadzu) equipped with a C18 column (Shimadzu), following the method described by Hamada and Maruta (2024).

Threonate extraction was performed according to Miyagi et al. (2010) with minor modifications. *Arabidopsis* shoots were frozen in liquid nitrogen and ground to a fine powder. Approximately 50 mg of the powdered sample was transferred to a pre-chilled tube and homogenized in 150 µL of 100% (v/v) methanol. Then, 150 µL of an internal standard solution containing 100 µM 1,4-piperazine diethane sulfonic acid (PIPES), 100 µM methionine sulfone (MeS), and 100 mM HCl was added. The homogenate was centrifuged at 15,300 × *g* for 5 min at 4°C. The supernatant was then filtered using a 3-kDa cutoff filter unit (Millipore) by centrifugation at 15,300 × *g* for 30 min. The resulting filtrate was analyzed by capillary electrophoresis–triple quadrupole mass spectrometry (CE-QQQ-MS) as described previously (Miyagi et al., 2019).

### Enzyme assays

Approximately 0.1 g of *Arabidopsis* shoots were frozen in liquid nitrogen and ground to a fine powder. The powder was homogenized in 1 mL of extraction buffer consisting of 50 mM Tris-HCl (pH 7.2) and 1 mM PMSF. The homogenate was centrifuged at 15,300 × *g* for 15 min at 4°C. The resulting supernatant was concentrated approximately 600-fold using an Amicon Ultra-4 centrifugal filter unit with a 30 kDa cutoff (PLBC Ultracel, Merck). The concentrated supernatant was used as the crude enzyme preparation.

LtnD activity staining was performed following the method described by Corpas et al. (2017), with modifications. Briefly, concentrated enzyme extracts were separated by 3–14% gradient native PAGE (ATTO) and incubated in a reaction mixture containing 0.8 mM NAD^+^, 10 mM calcium L-threonate, 0.24 µM NBT, and 65 nM PMS.

Spectrophotometric measurements of LtnD activity were performed using a microplate reader. The reaction solution consisted of 100 mM Tris-HCl (pH 8.0) containing 0.015 mM NBT, 0.003 mM PMS, 5% DMSO, and 1% Triton X-100. The substrate solution consisted of 100 mM Tris-HCl (pH 8.0) containing 1 mM NAD? and 1 mM calcium L-threonate. For each assay, 80 µL of the reaction solution and 10 µL of the substrate solution were pre-incubated at 37°C for 5 minutes. Then, 10 µL of the concentrated enzyme extract was added and mixed quickly. Formazan production was monitored by measuring absorbance at 570 nm. Because one molecule of calcium L-threonate contains two molecules of L-threonate, the final substrate concentration in the assay was 0.2 mM.

## Data Analyses

Statistical analysis methods are described in the figure legends (Dunnett’s test, or Student’s *t*-test). All calculations were performed using at least three independent biological replicates (see figure legends for details). In all experiments, shoots from at least ten plants were pooled and used as a single biological replicate.

## Data Availability Statement

The data underlying this article are available in the article and its online supplementary material.

## Acknowledgements

We thank Prof. Takahiro Ishikawa for valuable discussions and Mr. Naoya Odagaki for technical assistance.

## Author Contributions

K.Y. and T.M. conceived the project. K.Y., A.M., H.M., and T.M. designed the methodology. K.Y., A.M., and T.M. performed the formal analysis. K.Y., T.H., A.M., and A.H. conducted the investigation. K.Y., A.M., and T.M. curated the data. K.Y. and T.M. wrote the original draft. T.H., A.M., H.M., and A.H. contributed to reviewing and editing. T.M. prepared the visualizations. T.M. supervised the project and acquired funding.

## Supplementary materials

**Supplementary Table S1** List of primers used

**Supplemental Figure S1** Pairwise comparison of identity and similarity among LtnD-, OtnK-, and OtnC-like domains from *Arabidopsis* and bacterial enzymes

**Supplemental Figure S2** Domain architectures of LTD homologs in land plants

**Supplemental Figure S3** Purification of recombinant LtnD1-TF protein

**Supplemental Figure S4** Isolation and characterization of *ltd* mutant lines in Arabidopsis

**Supplemental Figure S5** Dehydrogenase activity toward various four-carbon organic acids in *Arabidopsis* wild type and *ltd-1*

