## Supplementary TableS1 for "Metabolism of L-threonate, an ascorbate degradation product, requires a protein with L-threonate metabolizing domains in *Arabidopsis*"

**Table S1** List of primers used

| Name | Sequence | Use |
|---|---|---|
| LTD-full-F | 5'-ATCATCATCATCATATGAGTGGCGTGGTTGGGT-3' | In-fution cloning |
| LTD-full-R | 5'-CGACAAGCTTGAATTTCATACCTTGATAAGGTTCTC-3' | In-fution cloning |
| LtnD1-R | 5'-TCATTGGCTAAGATCTCGTGAAATTCAAGCTTGTCG-3' | In-fution cloning |
| LtnD2-F | 5'-ATCATCATCATCATATGAGAATTGGTTTTATTGGT-3' | In-fution cloning |
| LtnD2-R | 5'-GGAGTGGTGAAGGTTTATTGAAATTCAAGCTTGTCG-3' | In-fution cloning |
| LTD-qF | 5'-TCCAGCTGAAGTGACGAAAGATG-3' | quantitative RT-PCR |
| LTD-qR | 5'-TCCTCAGCCTGAACCTCGTTTG-3' | quantitative RT-PCR |
| Actin2-qF3 | 5'-CTGTACGGTAACATTGTGCTCAG-3' | quantitative RT-PCR |
| Actin2-qR3 | 5'-CCGATCCAGACACTGTACTTCC-3' | quantitative RT-PCR |
| LTD-RT-F | 5’- TCTCGGAAAAAGTACTTGGTG-3’ | semi-quantitative RT-PCR |
| LTD-RT-R | 5’-ACCACTCCAGCATCATCTAT-3’ | semi-quantitative RT-PCR |
| Actin2-RT-F | 5’- GGCAAGTCATCACGATTGG -3’ | semi-quantitative RT-PCR |
| Actin2-RT-R | 5’- TCATACTCGGCCTTGGAGATC -3’ | semi-quantitative RT-PCR |
