## Supplementary Figures for "Metabolism of L-threonate, an ascorbate degradation product, requires a protein with L-threonate metabolizing domains in *Arabidopsis*"

Identiry

| LtnD | Arabidopsis-1 | Arabidopsis-2 | Escherichia | *Arthrobacter* |
|---|---|---|---|---|
| Arabidopsis-1 | 100 | 32.2 | 27.7 | 26.2 |
| Arabidopsis-2 | 32.2 | 100 | 38.7 | 36.8 |
| Escherichia | 27.7 | 38.7 | 100 | 34.1 |
| Arthrobacter | 26.2 | 36.8 | 34.1 | 100 |

| OtnK | Arabidopsis | Escherichia | Arthrobacter |
|---|---|---|---|
| Arabidopsis | 100 | 18.5 | 32 |
| Escherichia | 18.5 | 100 | 18.9 |
| Arthrobacter | 32 | 18.9 | 100 |

| OtnC | Arabidopsis | Escherichia | Arthrobacter |
|---|---|---|---|
| Arabidopsis | 100 | 1.5 | 33.2 |
| Escherichia | 1.5 | 100 | 22.6 |
| Arthrobacter | 33.2 | 22.6 | 100 |

Similarity

| LtnD | Arabidopsis-1 | Arabidopsis-2 | Escherichia | *Arthrobacter* |
|---|---|---|---|---|
| Arabidopsis-1 | 100 | 59.4 | 52.3 | 44.6 |
| Arabidopsis-2 | 59.4 | 100 | 60.7 | 49.7 |
| Escherichia | 52.3 | 60.7 | 100 | 52.6 |
| Arthrobacter | 44.6 | 49.7 | 52.6 | 100 |

| OtnK | Arabidopsis | Escherichia | Arthrobacter |
|---|---|---|---|
| Arabidopsis | 100 | 32.7 | 46.7 |
| Escherichia | 32.7 | 100 | 30.1 |
| Arthrobacter | 46.7 | 30.1 | 100 |

| OtnC | Arabidopsis | Escherichia | Arthrobacter |
|---|---|---|---|
| Arabidopsis | 100 | 2.8 | 47.7 |
| Escherichia | 2.8 | 100 | 22.6 |
| Arthrobacter | 47.7 | 22.6 | 100 |

| Color Scale |
|---|
| 80 |
| 60 |
| 40 |
| 20 |
| 0 |

**Supplementary Figure S1** Pairwise comparison of identity and similarity among LtnD-, OtnK-, and OtnC-like domains from *Arabidopsis* and bacterial enzymes

Heatmaps showing pairwise amino acid identity (left) and similarity (right) among representative LtnD-, OtnK-, and OtnC-like domains from Arabidopsis thaliana (AT1G18270.4), Escherichia coli (four-step pathway), and Arthrobacter sp. ZBG10 (three-step pathway). Two LtnD-like domains from Arabidopsis (LtnD1 and LtnD2) were included separately. Values represent pairwise percent identity or similarity based on global sequence alignment. Color intensity corresponds to the value according to the color scale shown on the right.

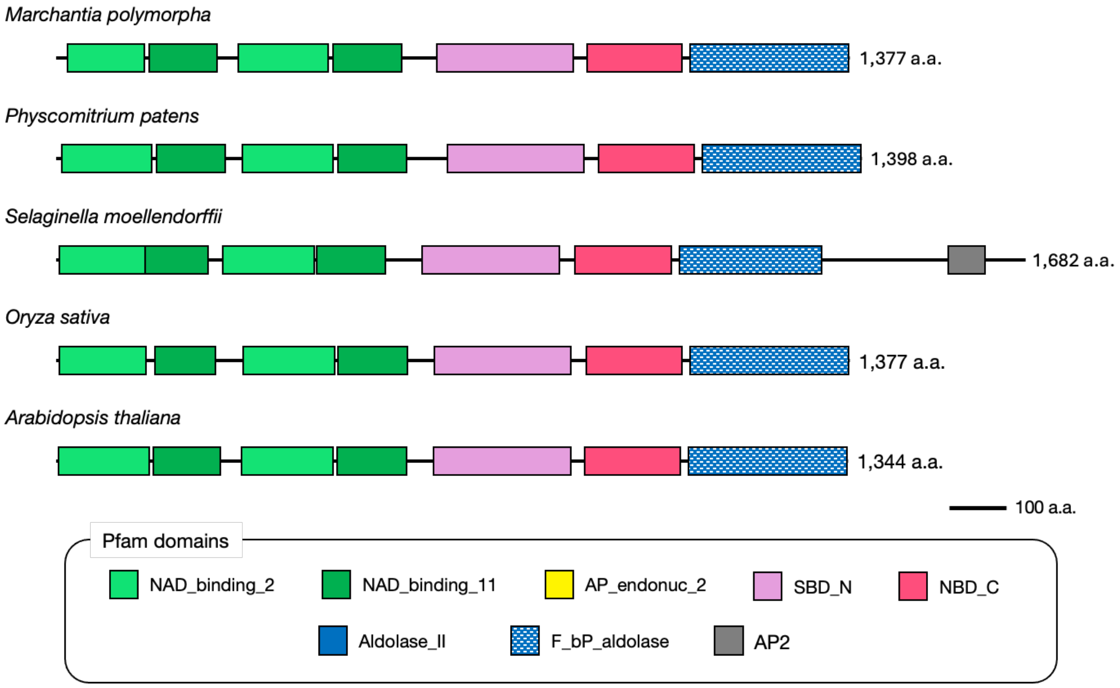


**Supplementary Figure S2** Domain architectures of LTD homologs in land plants

Schematic comparison of the domain structures of LTD homologs from representative land plant species: *Marchantia polymorpha*, *Physcomitrium patens*, *Selaginella moellendorffii*, *Oryza sativa*, and *Arabidopsis thaliana*. All homologs contain two LtnD-like domains (NAD_binding_2 and NAD_binding_11), one OtnK-like domain (SBD_N and NBD_C), and one OtnC-like domain (F_bP_aldolase), indicating a conserved domain architecture corresponding to the bacterial three-step L-threonate catabolic pathway. Interestingly, the *Selaginella* homolog contains an AP2 domain at the C-terminus, which is typically associated with transcriptional regulation. Domain annotations are based on Pfam predictions.

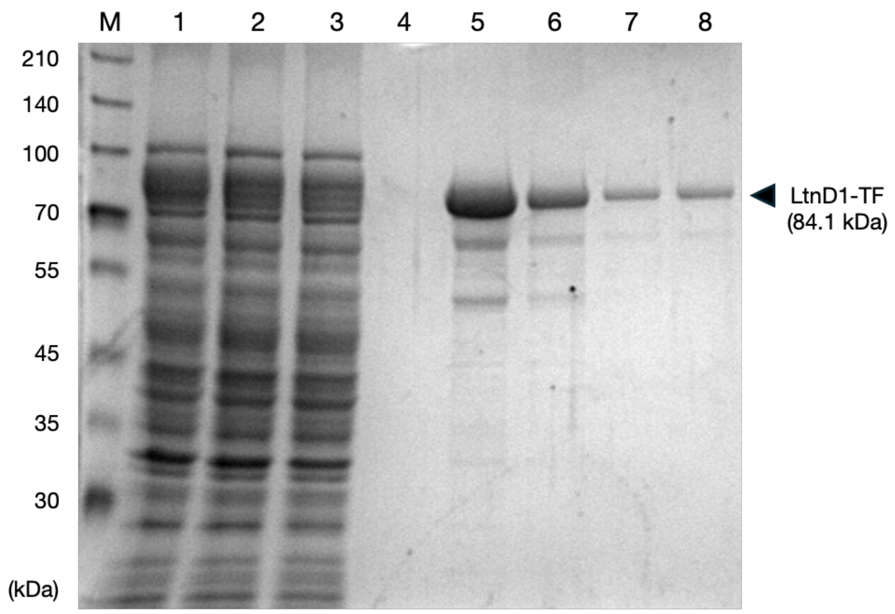


**Supplementary Figure S3** Purification of recombinant LtnD1-TF protein

Recombinant LtnD1 fused with a trigger factor (TF) tag (84.1 kDa) was expressed in *Escherichia coli* and purified using affinity chromatography. Protein samples were separated by SDS-PAGE and visualized by CBB staining. Lane M, molecular weight marker; lane 1, soluble fraction of crude extract; lane 2, flow-through from the affinity column; lanes 3 and 4, wash fractions (lane 3, first wash; lane 4, final wash); lanes 5–8, elution fractions. A prominent band corresponding to LtnD1-TF is visible in the eluted fractions (lanes 5–8), as indicated by the arrowhead. Eluted fractions from lanes 5–7 were used for activity assays; however, no detectable LtnD activity was observed.

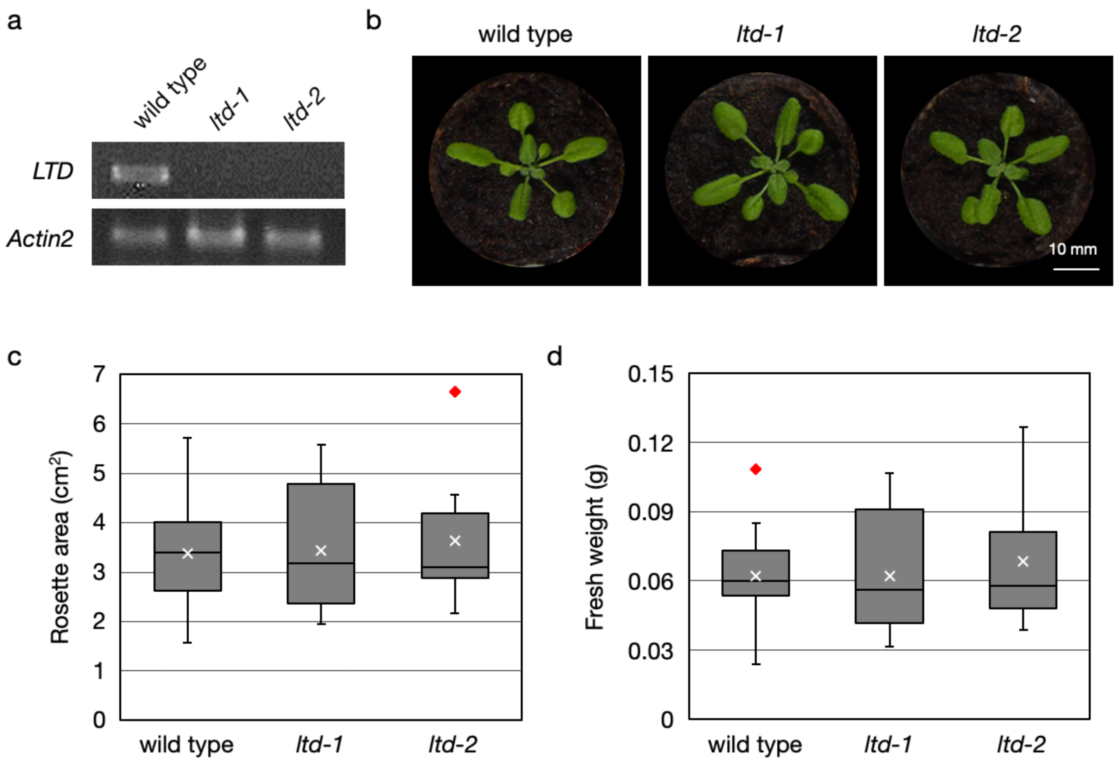


**Supplementary Figure S4** Isolation and characterization of *ltd* mutant lines in Arabidopsis

*Arabidopsis thaliana* wild-type (Col-0) plants were grown in soil for three weeks. (a) Semi-quantitative RT-PCR analysis of *LTD* transcript levels in wild-type, *ltd-1*, and *ltd-2* plants. *Actin2* was used as an internal control. (b) Representative images of 3-week-old plants grown under standard conditions. Scale bar: 10 mm. (c) Quantification of rosette area. (d) Shoot fresh weight. Each biological replicate included 5–6 plants, and measurements were performed on three independent biological replicates. Data from a total of more than 17 plants per genotype were used to generate box-and-whisker plots. In the plots, the center line represents the median, the box indicates the interquartile range, whiskers show the range, × marks the mean, and red diamonds indicate outliers. No statistically significant differences among genotypes were detected (one-way ANOVA).

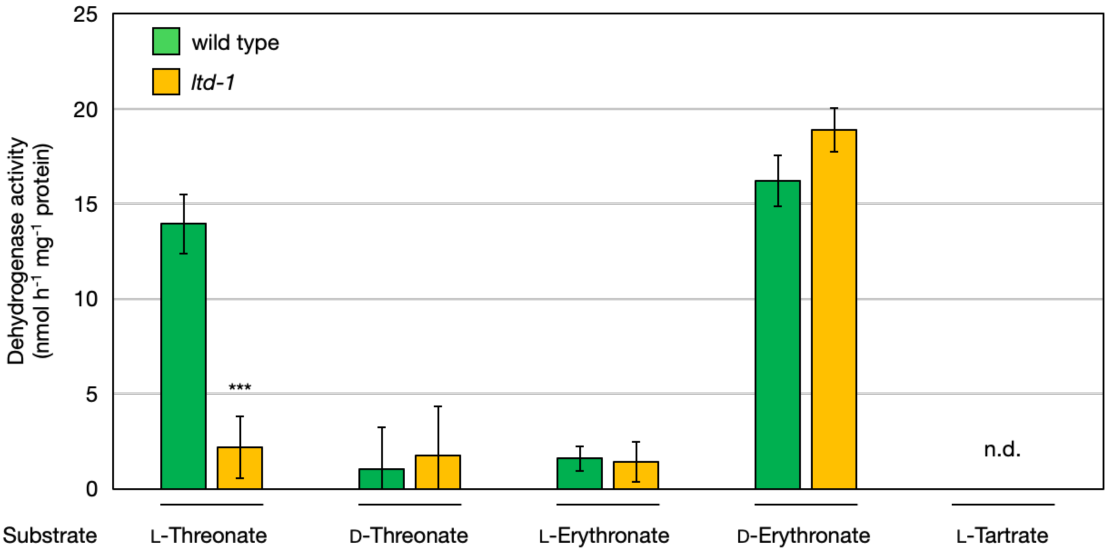


**Supplementary Figure S5** Dehydrogenase activity toward various four-carbon organic acids in *Arabidopsis* wild type and *ltd-1*

*Arabidopsis thaliana* wild-type (Col-0) and *ltd-1* plants were grown on half-strength MS medium without sucrose for two weeks. Shoots from 2-week-old plants were used to measure dehydrogenase activity toward various four-carbon organic acids. Substrates tested included L-threonate, D-threonate, L-erythronate, D-erythronate, and L-tartrate. Data are presented as mean ± SD of three biological replicates. Statistical differences between wild type and ltd-1 were assessed using Student's *t*-test. $^{***}P < 0.001$. n.d., not detected.
